# Choroid plexus APP regulates adult brain proliferation and animal behavior

**DOI:** 10.1101/734103

**Authors:** Karen Arnaud, Vanessa Oliveira Moreira, Jean Vincent, Glenn Dallerac, Chantal Le Poupon, Max Richter, Ulrike C. Müller, Laure Rondi-Reig, Alain Prochiantz, Ariel A. Di Nardo

## Abstract

Elevated amyloid precursor protein (APP) expression in the choroid plexus suggests an important role for extracellular APP metabolites in cerebrospinal fluid. Despite widespread *App* brain expression, we hypothesized that specifically targeting choroid plexus expression could alter animal physiology. Through various genetic and viral approaches in the adult mouse, we show that choroid plexus APP levels significantly impacted proliferation in both subventricular zone and hippocampus dentate gyrus neurogenic niches. Given the role of Aβ peptides in Alzheimer disease pathogenesis, we also tested whether favoring the production of Aβ in choroid plexus could negatively affect niche functions. After AAV5-mediated long-term expression of human mutated *APP* specifically in the choroid plexus of adult wild type mice, we observe reduced niche proliferation, behavioral defects in reversal learning, and deficits in hippocampal long-term potentiation. Our findings highlight the unique role played by the choroid plexus in regulating brain function, and suggest that targeting APP in choroid plexus may provide a means to improve hippocampus function and alleviate disease-related burdens.

## Introduction

The cerebrospinal fluid (CSF) has gained attention as a source of biomarkers mirroring the progression of Alzheimer disease (AD), the major cause of dementia worldwide (Henstridge *et al*, 2019). A clinically accepted CSF biomarker is the amount of Aβ42 relative to Aβ40 (Palmqvist *et al*, 2019; Molinuevo *et al*, 2018), both of which are peptide metabolites of the amyloid precursor protein (APP). As Aβ42 oligomerizes and forms amyloid plaques in the parenchyma, an inverse amount of soluble Aβ42 is cleared into the CSF. Along with the Aβ peptides, the soluble sAPPα and sAPPβ ectodomains have also been detected in CSF and were assumed to originate from the parenchyma through clearance (Perneczky *et al*, 2014). However, recent transcriptomics studies point to high APP expression in the choroid plexus (ChPl) (Liu *et al*, 2013; Baruch *et al*, 2014), a structure responsible for CSF secretion within brain ventricles. These findings suggest that APP metabolites from ChPl have functional purpose in the CSF, and we hypothesized that targeting ChPl expression of APP metabolites could alter animal physiology.

The ChPl has been recently considered an important contributor to adult neurogenesis by secreting trophic factors, transcription factors, and guidance molecules into the CSF which signal to the neurogenic niche (Lehtinen *et al*, 2013; Silva-Vargas *et al*, 2013; Bjornsson *et al*, 2015; Planques *et al*, 2019). Interestingly, a deficit in adult neurogenesis has been associated with AD (Moreno-Jiménez *et al*, 2019; Toda *et al*, 2019; Choi *et al*, 2018; Winner & Winkler, 2015; Rodríguez & Verkhratsky, 2011). Although adult neurogenesis in humans has been recently challenged (Sorrells *et al*, 2018) and debated (Paredes *et al*, 2018; Tartt *et al*, 2018), it has been demonstrated conclusively in rodents, with two primary neurogenic sites, the subventricular zone (SVZ) and the subgranular zone (SGZ) of the hippocampal dentate gyrus (DG). Neurogenesis has been shown to be impaired in mouse models of familial AD (FAD). Early models with mutated APP were found to have decreased adult SGZ proliferation and survival of neural progenitor cells (NPCs) (Haughey *et al*, 2002b; Donovan *et al*, 2006), while other models expressing FAD forms of APP and presenilin-1 were found to also show decreased adult SVZ proliferation (Zeng *et al*, 2016; Demars *et al*, 2010). Furthermore, deficits in adult neurogenesis were found to precede amyloid plaque and neurofibrillary tangle formation (Hamilton *et al*, 2010).

APP has been shown to play a role in adult neurogenesis, either as full-length transmembrane APP (Wang *et al*, 2014) or through its sAPPα ectodomain (Demars *et al*, 2011; Caillé *et al*, 2004; Baratchi *et al*, 2012). While the controversy on adult neurogenesis in the humans cannot be ignored, recent studies convincingly show that adult DG neurogenesis is maintained at a high level until old age but drops sharply in AD patients (Moreno-Jiménez *et al*, 2019) and in patients with mild cognitive impairments (Tobin *et al*, 2019), supporting the AD “neurogenesis” hypothesis of cognitive impairment (Choi & Tanzi, 2019; Lazarov & Hollands, 2016; Lazarov *et al*, 2010). We previously showed that sAPPα infused in the CSF recognizes stem cells expressing the EGF-receptor within the SVZ and enhances their proliferation, while no effect on hippocampal neurogenesis was observed (Caillé *et al*, 2004). In line with these observations, we aimed to evaluate the importance of ChPl APP in regulating adult neurogenesis and to explore the possibility that the ChPl might constitute an important actor and translational target in healthy and pathological aging. We report that sAPPα produced specifically from the ChPl positively affects proliferation in both neurogenic niches. Conversely, ChPl-specific viral expression of human APP bearing the Swedish-Indiana (SwInd) mutations to favor CSF Aβ production, resulted in reduced proliferation in both niches, caused defects in reversal learning and impaired synaptic plasticity mechanisms in the hippocampus.

## Results

### App is highly expressed in the adult choroid plexus

Given that ventricular infusion of recombinant sAPPα has been shown to affect SVZ proliferation (Caillé *et al*, 2004; Demars *et al*, 2013), and that the ChPl has been shown to express APP (Liu *et al*, 2013), we hypothesized that the ChPl is a potential endogenous source for CSF-borne sAPPα. Indeed, *App* is one of the most highly expressed ChPl genes (Baruch *et al*, 2014). We confirmed high *App* expression levels in the ChPl by qPCR. In comparison to the hippocampus (that has been shown previously to strongly express APP) and the SVZ, we found higher and elevated levels in ChPl in lateral and 4th ventricles of 4-month-old adult mice (Fig. 1*A*). Interestingly, *Transthyretin* and *Apoe*, two genes involved in App functions (Sousa *et al*, 2007; Deane *et al*, 2008) are also very highly expressed in the ChPl (Baruch *et al*, 2014; Silva-Vargas *et al*, 2016).

**Figure 1.**
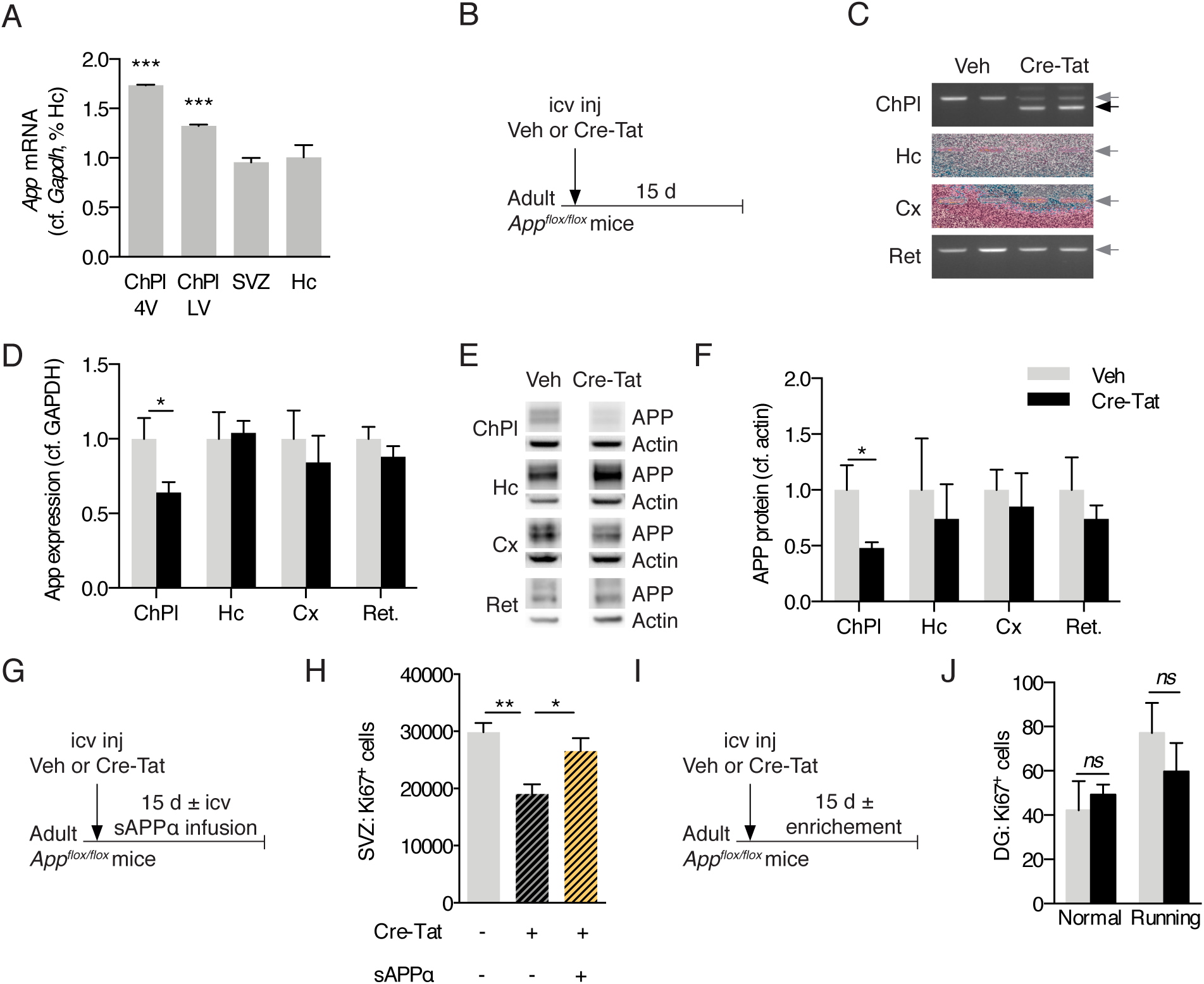
Choroid plexus App loss-of-function in *App^flox/flox^* mice decreases adult neural progenitor proliferation. **A** Quantitative PCR analysis of *App* expression in ChPl from ventricles (LV and 4V), SVZ, and Hc. Values are normalized to GAPDH and expression in Hc (Veh n = 4, Cre-Tat n=7). **B** Schematic of App ChPl knock-down model involving single icv injection of Veh or Cre-Tat in adult *App^flox/flox^* mice. **C** *App* specific recombination in LV choroid plexus after Cre-Tat icv injection. PCR of genomic DNA extracted from indicated structures. Grey arrow indicates *App* wild type locus, black arrow indicates deleted *App* locus upon Cre recombination. **D** Quantitative PCR analysis of *App* expression normalized to GAPDH after Cre-Tat or Veh icv injection (Veh n = 10, Cre-Tat n=10). **E-F** Western blot analysis (**E**) of various brain structures after Cre-Tat or Veh icv injection for quantification (**F**) of APP protein levels normalized to actin (Veh n = 10, Cre-Tat n=10). **G** Schematic of sAPPα rescue paradigm for evaluating SVZ cell proliferation. sAPPα was infused by 15-day osmotic mini-pump implanted immediately after Cre-Tat icv injection. **H** Analysis of SVZ cell proliferation by quantification of Ki67 positive cells after Veh or Cre-Tat icv injection followed by infusion of Veh or sAPPα (Veh/Veh n=7, Cre-Tat/Veh n= 6, Cre-Tat/sAPPα n=7). **I** Schematic of App ChPl knock-down with environmental enrichment (running wheel) for evaluating DG cell proliferation. **J** Analysis of DG cell proliferation by quantification of Ki67 positive cells after *App* recombination in mice in normal conditions or after physical exercise (Veh n=7, Cre-Tat n=8). *p<0.05; **p<0.01; *t* test in **D**, **F**, and **J**; one-way ANOVA in **H**; all values, mean ±SEM. LC, lateral ventricle; 4V, 4^th^ ventricle; ChPl, choroid plexus; Hc, hippocampus; Cx, cortex; Ret, retina; SVZ, subventricular zone; DG, dentate gyrus; icv, intracerebroventricular; Veh, vehicle.

### App knock-down in the choroid plexus decreases adult proliferation

In order to knock down *App* expression locally and specifically in ChPl, we performed Cre-Tat intracerebroventricular (icv) injections in 10-month-old *App^flox/flox^* mice (Fig. 1*B*). Cre-mediated deletion of the *App* locus and *App* expression were evaluated 15 days later in various structures including retina, hippocampus, cortex and ChPl (Fig. 1*B-F*). The specificity of Cre-Tat targeting selectively the ChPl was confirmed by the absence of recombination in other structures (Fig. 1*C*), as previously demonstrated for another floxed mouse model (Spatazza *et al*, 2013). Consequently, only in ChPl do we observe a decrease in *App* mRNA (by 37 ± 7%, Fig. 1*D*) and APP protein (by 52 ± 3%, Fig. 1*E-F*).

To evaluate the impact of *App* knock-down on cell proliferation, ~3-month-old *App^flox/flox^* mice were injected with vehicle or Cre-Tat and subsequently implanted with 15-day osmotic minipumps for CSF infusion of either vehicle or sAPPα (Fig. 1*G*). Compared to vehicle injected/infused controls, animals injected with Cre-Tat and infused with vehicle showed a significant reduction in the number of proliferating cells in the SVZ (Fig. 1*H*). This decrease was rescued by infusion of sAPPα (Fig. 1*H*), suggesting that knock-down of sAPPα leads to impaired neurogenesis. In contrast, this experimental paradigm did not alter DG proliferation (Fig. 1*I-J*), even under enriched environment conditions with free access to running wheels known to stimulate hippocampal proliferation (Praag *et al*, 1999).

In order to reduce *App* levels more robustly in the ChPl, a viral vector expressing shRNA against the mouse *App* sequence was injected into the ventricles of ~3-month-old wild type mice (Fig. 2*A*). As previously reported (Watson *et al*, 2005), icv injection of serotype 5 adeno-associated virus (AAV5) results in specific ChPl targeting, and indeed we did not detect expression of co-expressed eGFP marker protein in the parenchyma (Fig. 2*B*). Eight weeks post-injection, *App* (mRNA) and APP protein decreased nearly 5-fold in ChPl with no change in hippocampus and cortex levels (Fig. 2*C-E*). This decrease led to a significant reduction in the number of proliferating cells in both SVZ and DG (Fig. 2*F-I*). Thus, there appear to be differences in sensitivity between SVZ and DG: in the *App^flox/flox^* mouse model, where ChPl *App* (mRNA) knockdown is ~50%, proliferation decreases in SVZ but not in DG; in the viral shRNA(*App*) model, where ChPl *App* (mRNA) knockdown is ~80%, proliferation decreases in both niches.

**Figure 2.**
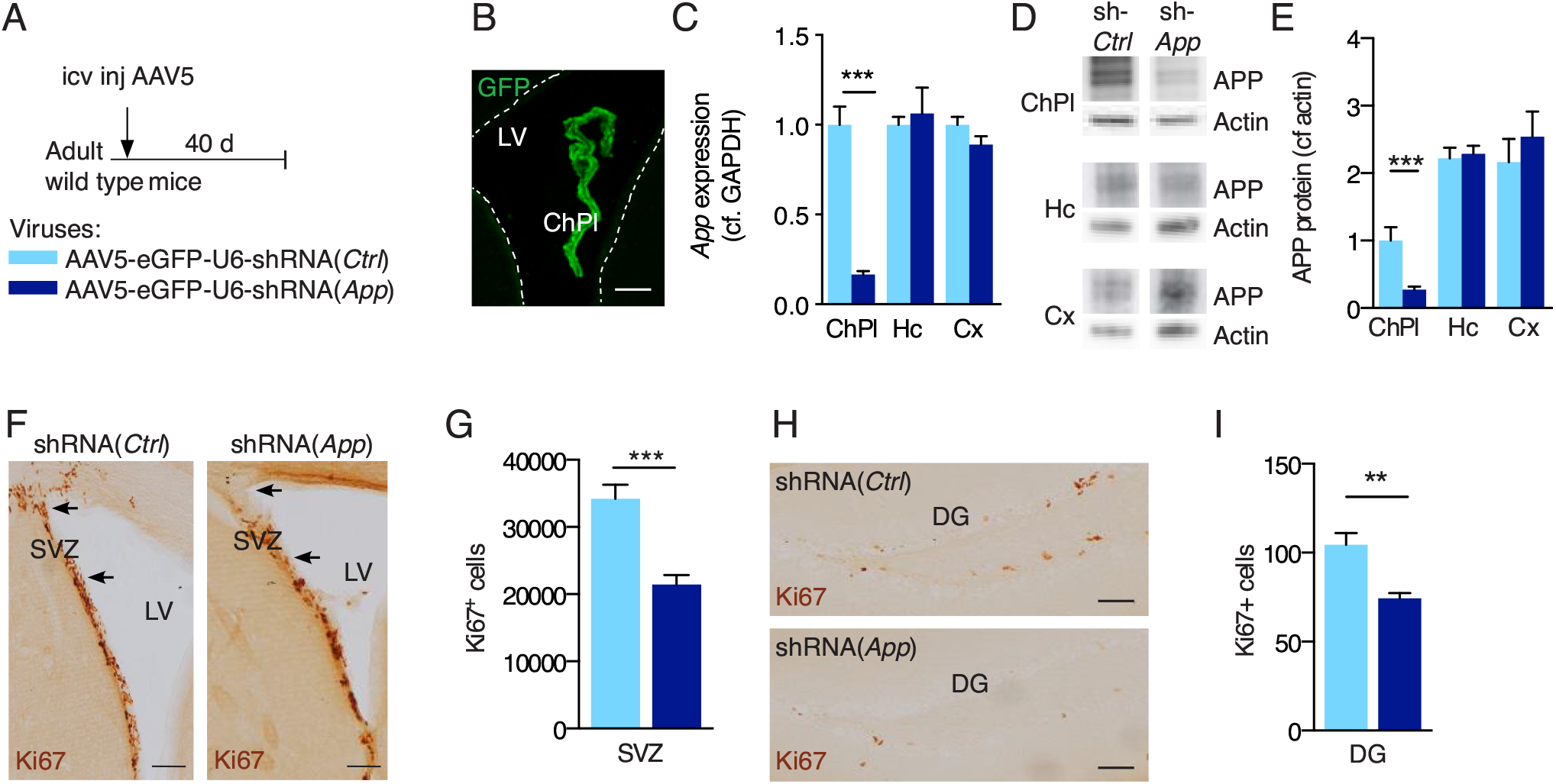
Knock-down of choroid plexus App expression reduces cell proliferation in neurogenic niches. **A** Schematic of App ChPl knock-down model involving a single icv injection of AAV5 expressing shRNA targeting the mouse *App* sequence in wild type mice. Control (Ctrl) virus expresses a scrambled shRNA devoid of targets in mouse. **B** An icv injection of AAV5 leads to specific choroid plexus expression. Note the absence of eGFP in brain parenchyma. Scale bar, 80 μm. **C** Quantitative PCR analysis of *App* expression in the ChPl, Hc, and Cx after ChPl *App* knockdown (shRNA-*App* n=4, shRNA-*Ctrl* n=4). **D-E** Western blot analysis (**D**) of various brain structures after ChPl *App* knock-down for quantification (**E**) of APP protein levels normalized to actin (shRNA-*App* n=4, shRNA-*Ctrl* n=4). **F-I** Analysis of cell proliferation in SVZ (**F**,**G**) and DG (**H**,**I**) by quantification of Ki67 positive cells after ChPl *App* knock-down (shRNA-*App* n=7, shRNA-*Ctrl* n=5). Arrows (**F**) highlight regional differences in SVZ cells. Scale bars, 100 μm. **p<0.01; ***p<0.001; *t* test; all values, mean ± SEM. ChPl, choroid plexus; Hc, hippocampus; Cx, cortex; SVZ, subventricular zone; DG, dentate gyrus; icv, intracerebroventricular.

### sAPPα gain-of-function in either cerebrospinal fluid or choroid plexus increases adult proliferation

APP can be cleaved either along the non-amyloidogenic pathway to give rise to sAPPα or along the amyloidogenic pathway to produce its sAPPβ counterpart, and the icv injection of the two sAPP forms has been previously shown to have opposite effects on neurogenesis. sAPPα was found to increase both SVZ and SGZ NPC proliferation 30 hours after icv injection of sAPPα protein in ~8-month-old wild type mice, while sAPPβ protein icv injection decreased proliferation in both niches (Demars *et al*, 2013). However, a previous study involving 3-day icv infusion of sAPPα-Fc in ~2-month-old wild type mice found increased SVZ proliferation but no change in SGZ proliferation (Caillé *et al*, 2004); these differences may be due to longer infusion times or to the use of a fusion sAPPα-human IgG1(Fc) protein construct. To further investigate the impact of sAPP on adult neurogenesis, either sAPPα or sAPPβ protein was infused for 7 days in the lateral ventricles of ~3-month-old wild type mice (Fig. 3*A*). In our infusion paradigm, sAPPα protein increased the number of proliferating cells both in the SVZ and SGZ of adult mice, while sAPPβ protein had no effect (Fig. 3*B-E*). To confirm that CSF and choroid plexus sAPPα gain-of-function are correlated, icv injections of AAV5 expressing mouse *App* were performed for ChPl-specific over-expression (Fig. 3*F*). After 8 weeks, the relative *App* expression was increased by approximately 2-fold (Fig. 3*G*), resulting in a significant increase in the number of proliferating cells in both the SVZ and SGZ (Fig. 3*H-I*).

**Figure 3.**
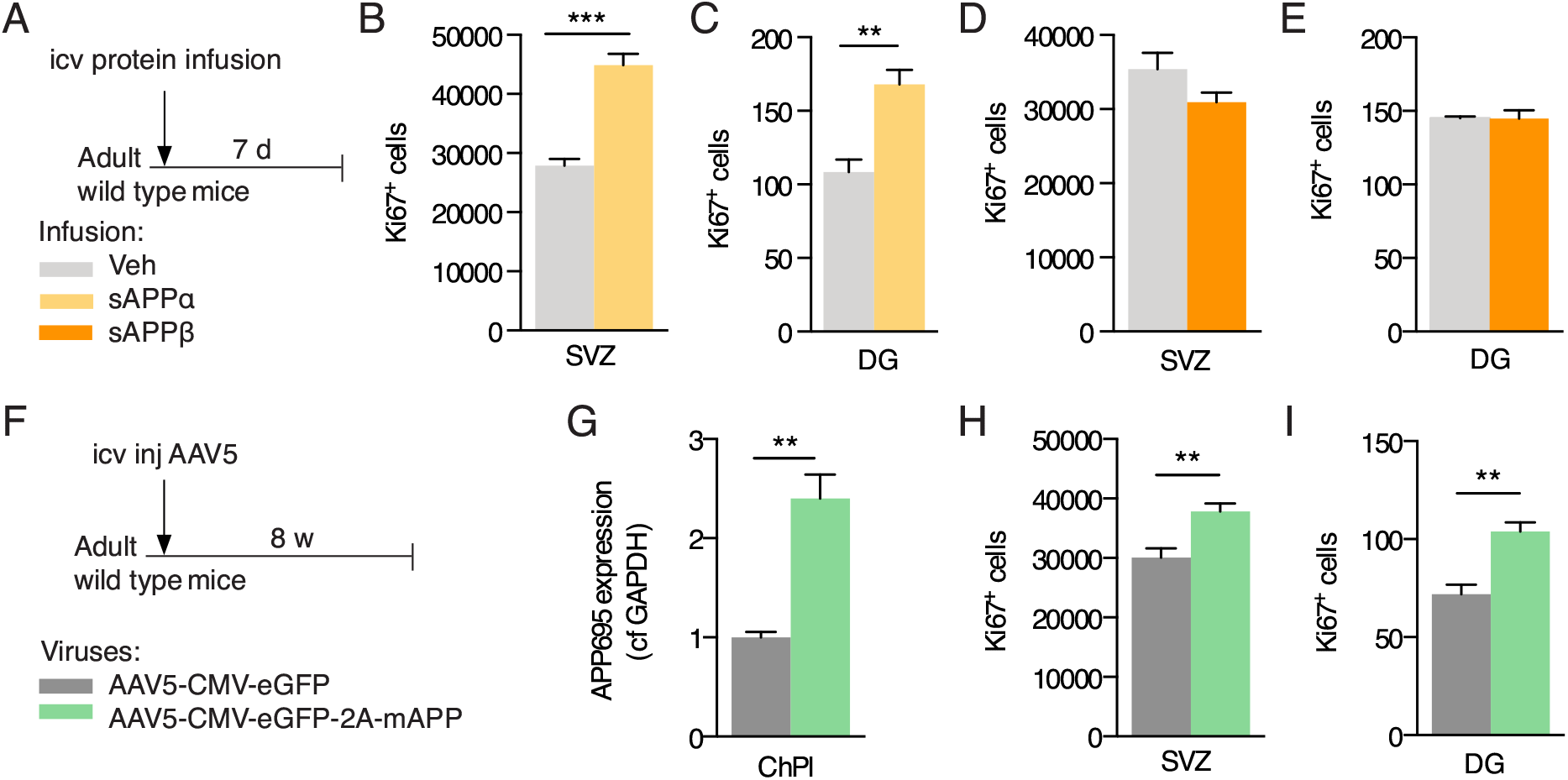
sAPPα gain-of-function in the cerebral ventricles increases cell proliferation in neurogenic niches. **A** Schematic of sAPP icv infusion by 7-day osmotic mini-pump in adult wild type mice. **B-C** Analysis of cell proliferation in SVZ (**B**) and DG (**C**) by quantification of Ki67 positive cells after icv infusion of sAPPα or Veh (Veh n=9, sAPPα n=9). **D-E** Analysis of cell proliferation in SVZ (**D**) and DG (**E**) by quantification of Ki67 positive cells after icv infusion of sAPPβ or Veh (Veh n=5, sAPPβ n=5). **F** Schematic of App ChPl gain-of-function model involving a single icv injection of AAV5 expressing mouse *App* or eGFP (control) in wild type mice. **G** Quantitative PCR analysis of *App* expression in ChPl after AAV5 icv injection (AAV5-mAPP n=4, AAV5-eGFP n=3). **H-I** Analysis of cell proliferation in SVZ (**H**) and DG (**I**) by quantification of Ki67 positive cells after icv injection (AAV5-mAPP n=6, AAV5-eGFP n=5). *p<0.05; **p<0.01; ***p<0.001; *t* test; all values, mean ± SEM. ChPl, choroid plexus; Hc, hippocampus; SVZ, subventricular zone; DG, dentate gyrus; icv, intracerebroventricular; Veh, vehicle.

### Choroid plexus expression of APP(SwInd) impairs behavior

Because altering APP expression selectively in the ChPl impacted neurogenic niches, we hypothesized that favoring the production of Aβ specifically in ChPl could negatively affect niche functions. To explore this possibility, we overexpressed a mutated form of human APP (Swedish K670N/M671L and Indiana V17F mutations) specifically in the ChPl of wild type mice at 3 months of age (Fig. 4*A*). The consequences of this gain-of-function were evaluated 3 and 12 months after injection (when mice were 6 and 15 months old, respectively). We found strong *hAPP(SwInd)* mRNA expression in ChPl after 3 months which increased more than 3-fold by 12 months (Fig. 4*B*). Strikingly, proliferation was significantly decreased at 3 months post-injection in both SVZ and DG of hAPP(SwInd) expressing mice. However, no difference was seen at 12 months post-injection (Fig. 4*C-D*), although this is likely due to the extremely low levels of proliferation also observed in control mice that may preclude further reduction. Indeed, proliferation already dramatically decreases from the time of injection at 3 months (‘No injection’ in Fig. 4*C-D*) to 6 months (3 months post-injection) in both niches, and further decreases in the DG at 15 months (12 months post-injection) as previously reported (Encinas *et al*, 2011). This decrease in proliferation was not due to amyloid plaque accumulation, as none was observed in the cortex or hippocampus of mice expressing the human *APP(SwInd)* in the ChPl (data not shown).

**Figure 4.**
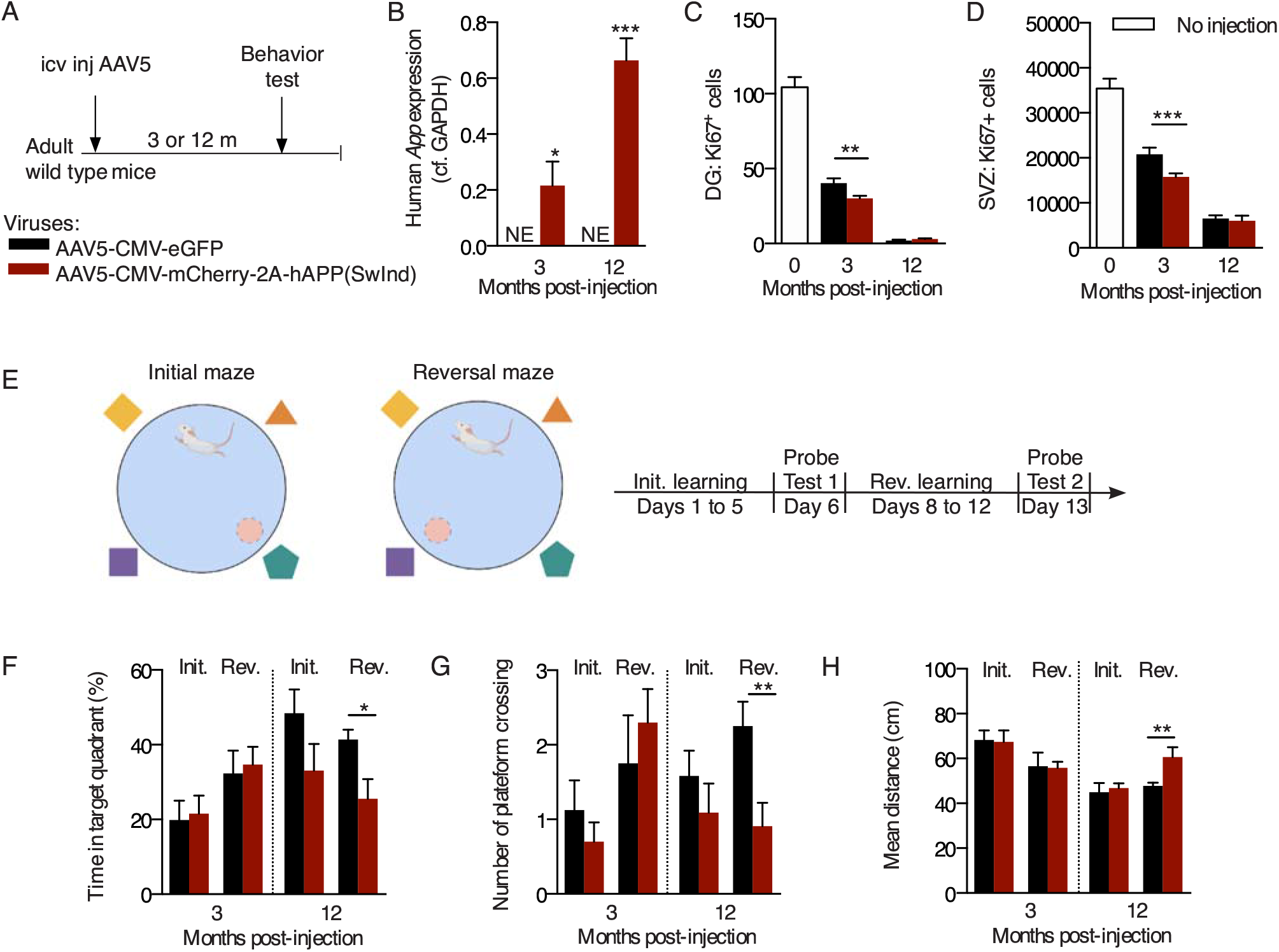
Choroid plexus hAPP(SwInd) expression decreases proliferation in neurogenic niches and impairs reversal learning. **A** Schematic of mutant APP ChPl model involving a single icv injection of AAV5 expressing human mutated *APP* or eGFP (control) in wild type mice. **B** Quantitative PCR analysis of ChPl viral expression of hAPP(SwInd) in WT mice at 3- and 12-months post-injection (n=6 per group). **C-D** Quantification of Ki67 positive cells in SVZ (**C**) and DG (**D**) after ChPl viral expression (AAV5-eGFP n=6, AAV5-hAPP(SwInd) n=12). **E** Reversal learning paradigm. **F-H** Behavioral responses after probe test 1 (Initial) and probe test 2 (Reversal) quantified by time (**F**), activity (**G**) and distance (**H**) (6 months: AAV5-eGFP n=9, AAV5-hAPP(SwInd) n=11; 15 months: AAV5-eGFP n=11, AAV5-hAPP(SwInd) n=10). *p<0.05; **p<0.01; ***p<0.001; *t* test; all values: mean ± SEM. SVZ, subventricular zone; DG, dentate gyrus; NE, not expressed; icv, intracerebroventricular.

We also evaluated spatial memory by using the Morris water maze reversal learning paradigm at either 3 months or 12 months post-injection (Fig. 4*E*). Mice from either group had not been subjected to previous spatial memory tests. During the learning phases, all groups showed significant improvement in latency to find the hidden platform after 5 days of training, but no differences were observed between groups (control and *hAPP(SwInd)*) in either latency or swimming speed at both ages (Supp. Fig. 1). During probe tests, platforms were removed and performance was evaluated by the time spent in the target platform quadrant, the number of platform area crossings, and the mean total distance taken to reach the platform area (Fig. 4*F-H*). Swimming speed remained unaltered during probe tests (Supp. Fig. S1). At either 3 months or 12 months post-injection, *hAPP(SwInd)* mice showed no difference in performance compared to controls during probe test 1 (initial learning). Performance after probe test 2 (reversal learning) was unchanged in 6-month-old mice (3 months post-injection) but showed significant differences for all parameters at 15 months (12 months post-injection). The time spent in the trained target quadrant was decreased, as was the number of platform area crossings, while the mean distance to find the platform area was increased, indicating that reversal learning is impaired after long-term ChPl expression of *hAPP(SwInd)*.

### Choroid plexus expression of App(SwInd) impairs synaptic plasticity

Compromised reversal learning may be rooted in plasticity-dependent deficits in hippocampal-dependent spatial memory. Interestingly, mouse models of AD show a marked decrease in hippocampal CA1 synaptic plasticity in the form of long-term potentiation (LTP) at 4 months of age (Li *et al*, 2017). We thus assessed synaptic function at Schaffer collateral CA3 to CA1 synapses in the hippocampus of 15-month old mice expressing *hAPP(SwInd)* in the ChPl after receiving AAV5 icv injections at 3 months of age (Fig. 5*A*). We found no change in basal synaptic transmission in CA1 as shown by comparable input/output curves (F>1) (Fig. 5*B*). However, paired-pulse facilitation, a form of short-term plasticity reflecting presynaptic function, is increased in mice expressing mutated APP thus indicating a decrease in pre-synaptic release probability (Fig. 5*C*). Consistently, induction of LTP by high-frequency stimulation is significantly impaired in *AAV5-hAPP(SwInd)* injected mice (Fig. 5*D*), revealing impaired plasticity which corroborates observed deficits in spatial memory (Fig. 4*F-H*).

**Figure 5.**
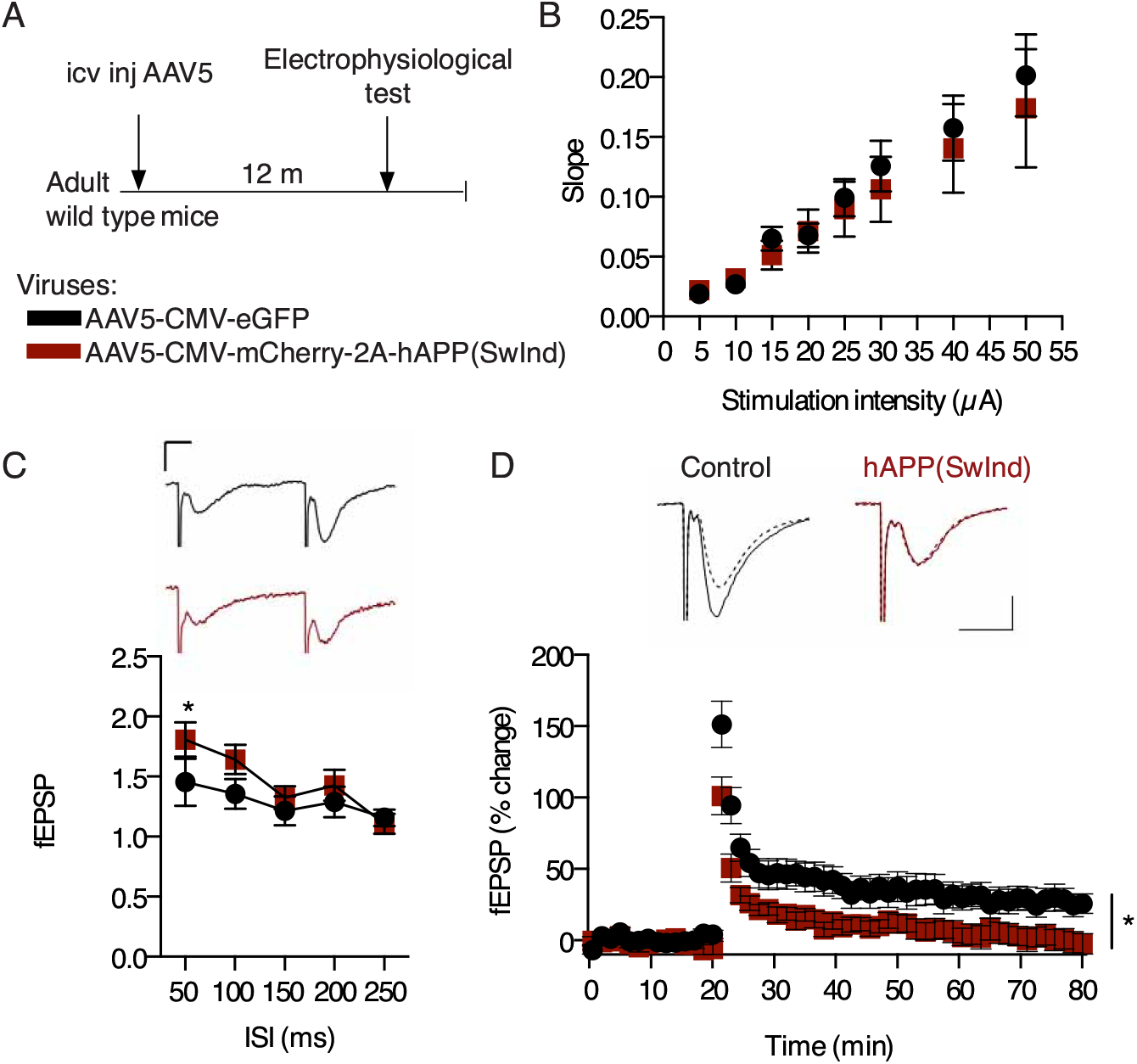
Choroid plexus hAPP(SwInd) expression impairs hippocampal LTP. **A** Schematic of mutant APP ChPl model involving a single icv injection of AAV5 expressing human mutated *APP(SwInd)* or eGFP (control) in wild type mice. **B** Comparaison of stimulus-response (input/output) relationship in CA1 region between AAV5-hAPP(SwInd) (*N*=5, *n*=13, black squares) and AAV5-eGFP (*N*=7, *n*=10, red dots) injected mice. **C** Comparison of paired pulse facilitation between AAV5-hAPP(SwInd) (*N*=6, *n*=13) and AAV5-eGFP (*N*=8, *n*=13) injected mice. **D** LTP was induced by high frequency stimulation (HFS) at Schaffer collateral-CA1 synapses after 20 min baseline recording of slices from mice injected with AAV5-hAPP(SwInd) (*N*=4, *n*=9) or AAV5-eGFP (*N*=5, *n*=7). Representative traces showing responses before (dashed line) and 60 min after tetanus delivery (bold line). *p<0.05; ANOVA; all values: mean ± SEM; *N*, number of animals; *n*, number of slices; icv, intracerebroventricular. Calibration bars: 10 ms, 0.2 mV.

## Discussion

The present study takes its origin in the growing interest for the ChPl, a structure increasingly recognized for its physiological importance beyond its classical “kidney of the brain” functions as it secretes CSF containing a plethora of trophic compounds (Ghersi-Egea *et al*, 2018). Among these compounds are growth factors, guidance cues and morphogens, some of which gain access to the SVZ and participate in the regulation of neuroblast migration (Planques *et al*, 2019; Silva-Vargas *et al*, 2016; Falcão *et al*, 2012). sAPPα infused into the CSF has been shown to increase proliferation and progenitor cell numbers in the SVZ (Demars *et al*, 2013; Caillé *et al*, 2004), with binding sites on type C and type A progenitor cells (Caillé *et al*, 2004). Given the very high expression of APP by the ChPl, we hypothesized that sAPPα could be secreted by the ChPl into the CSF and participate in adult neurogenesis. By conditionally knocking down *App* expression specifically in the ChPl through either genetic deletion or shRNA expression, we establish that ChPl APP has neurogenic activity not only in the SVZ but surprisingly also in the DG. Furthermore, this reduced proliferation can be reversed by the infusion of sAPPα into the CSF. Thus, sAPPα secreted by the ChPl can not only gain access to cells within the “superficial” SVZ, which contact lateral ventricles, but also to cells within the “deeper” DG within the parenchyma. Such transport has been reported for transcription factors secreted into the CSF by the ChPl (Spatazza *et al*, 2013). Together, these results strongly suggest that sAPPα secreted by the ChPl functions as a neurogenic factor, further confirming ChPl as a major actor in regulating adult neurogenesis.

The full range of functions of the APP family have yet to be fully identified, but several studies strongly suggest important developmental and physiological roles (Müller *et al*, 2017). Key functions of sAPPα include its ability to enhance spine density, synaptic plasticity, and memory (Mockett *et al*, 2019; Müller *et al*, 2017). Expression of sAPPα in hippocampus has been shown to rescue cognition and synaptic transmission and to mitigate synaptic and cognitive deficits in a pathological context (Fol *et al*, 2016; Xiong *et al*, 2017; Tan *et al*, 2018; Rice *et al*, 2019). Certain functions involve *cis* or *trans* dimer interactions between transmembrane APP molecules, and others involve heterodimers between transmembrane APP and secreted sAPPα (Milosch *et al*, 2014). Some of these properties are attributed to the extracellular C-terminal 16 amino acids of sAPPα, as compared to sAPPβ. For example, acute *in vitro* or *in vivo* application of sAPPα but not sAPPβ rescues hippocampal LTP in the adult brain of conditional double knockout mice lacking APP and the related protein APLP2 (Hick *et al*, 2015; Richter *et al*, 2018). This C-terminal sequence facilitates synaptic plasticity in the hippocampus through binding to functional nicotinic α7-nAChRs (Richter *et al*, 2018; Morrissey *et al*, 2019). In keeping with previous results (Demars *et al*, 2013), we also find that *in vivo* icv infusion of sAPPα, but not sAPPβ, positively impacts neurogenic niche cell proliferation in the adult mouse brain. While the reason for this functional divergence between sAPPα and sAPPβ for proliferation remains unknown, a shift in APP processing towards the amyloidogenic pathway would clearly impair both synaptic plasticity and neurogenesis.

Mouse models developed to analyze the role of specific human *APP* mutations typically employ mutated genes often expressed throughout the brain and even the body. Our study is unique in that it explores whether expressing mutated APP only in the ChPl of adult mice (3 months old) could affect neurogenic niche proliferation and trigger learning defects. Specific expression in ChPl of human *APP(SwInd)* mutant, which favors sAPPβ and Aβ peptide formation, decreases proliferation at 3 months after infection (6 months old) with no change at 12 months after infection (15 months old). This lack of change is likely due to the naturally low level of neurogenesis at 15 months which leaves little room for significant further decrease. However, a reversal learning test showed a significant decrease in the cognitive performance of these mice at 15 months. Accordingly, electrophysiological analysis at 15 months revealed decreased pre-synaptic release probability and impaired synaptic plasticity. No amyloid plaques were observed in these mice, leading us to speculate that the functional outcome of increased sAPPβ and Aβ accumulation is rooted in either synaptic deficits, reduced neurogenesis, and/or neuronal aging (Suberbielle *et al*, 2013). Given that sAPPβ icv infusion does not affect proliferation in the neurogenic niches, it is more likely that the observed deficits are due to increased Aβ (soluble or oligomers). Indeed, icv injection of Aβ has been shown to decrease SVZ proliferation (Haughey *et al*, 2002a). Finally, while we observe no change in proliferation at 15 months, we cannot discount a negative long-term effect of ChPl human *APP(SwInd)* on the shaping of hippocampal synaptic circuits due to impaired proliferation during aging.

Our findings strengthen the unique role played by the ChPl in regulating adult neurogenesis, and it is important to consider the morphological and transcriptomic alterations of the ChPl during normal aging and in late-onset AD that can impact the brain (Silva-Vargas *et al*, 2016; Serot *et al*, 2001, 2000, 2012; Balusu *et al*, 2016). These alterations can lead to changes in blood-CSF barrier properties, in ChPl function regulation, as well as in CSF composition and turnover. Furthermore, adult neurogenesis failure clearly plays a role in the development of AD in familial and possibly sporadic AD patients (Mu & Gage, 2011; Rodríguez & Verkhratsky, 2011; Hamilton *et al*, 2013; Gonçalves *et al*, 2016; Moreno-Jiménez *et al*, 2019; Toda *et al*, 2019) and could be in part responsible for the decrease in neurogenesis observed in AD patients (Moreno-Jiménez *et al*, 2019). Indeed, cognition in a FAD mouse model can be improved by enhancing SGZ neurogenesis and elevating levels of brain-derived neurotrophic factor (Choi *et al*, 2018). From a translational viewpoint, the fact that expressing a mutated *APP* gene exclusively in the ChPl can alter the cognitive abilities of mice raises the possibility that modifying the expression of APP or targeting APP mutations specifically in the ChPl, a structure accessible from the venous compartment (Spatazza *et al*, 2013), may represent a novel means to alleviate the burden associated with AD.

## Materials and Methods

### Animals and ethics

C57Bl/6J mice were purchased from Janvier (France) and *App^flox/flox^* mice were described previously (Mallm *et al*, 2010). All colonies were maintained under a 12:12 light/dark cycle with free access to food and water. In environmental enrichment experiments, *App^flox/flox^* mice were placed in cages with running wheels (2 mice per cage) 7 days before surgery and kept for 15 days in these cages before analysis. All animal procedures were carried out in accordance with the guidelines of the European Economic Community (2010/63/UE) and the French National Committee (2013/118). For surgical procedures, animals were anesthetized with xylazine (Rompun 2%, 5 mg/kg) and ketamine (Imalgene 1000, 80 mg/kg) by intraperitoneal injection. This project (no. 00702.01) obtained approval from Ethics committee no. 59 of the French Ministry for Research and Higher Education.

### Protein and virus stereotaxic surgery

Vectored Cre recombinase protein, Cre-Tat, was produced as previously described (Spatazza *et al*, 2013). Adeno-associated virus (AAV) were of serotype 5 and purchased from either SignaGen [AAV5-CMV-eGFP-U6-shRNA(*App*) and AAV5-CMV-eGFP-U6-shRNA(*Ctrl*)] or Vector Biolabs [AAV5-CMV-eGFP-2A-mAPP[NM_007471.3], AAV5-CMV-mCherry-2A-hAPP(SwInd), and AAV5-CMV-eGFP]. The hAPP(SwInd) sequence is a form of human APP [NM_201414.2] bearing both Swedish (K670N/M671L) and Indiana (V717F) related mutations. Cre-Tat protein (~30 μg), vehicle (protein vehicle is 1.8% NaCl, 15% DMSO; virus vehicle is 0.9% NaCl), or high-titer AAVs (~10^13^ GC/ml) were injected (2 μl per mouse) into the right lateral ventricle (coordinates from bregma: x, −0.58 mm; y, +1.28 mm; z, −2 mm) with a 10 μl Hamilton syringe (0.2 μl/min). For protein infusion (3.5 μg per mouse), sAPPα (Sigma-Aldrich), sAPPβ (Sigma-Aldrich), or vehicle were infused for either 7 or 15 days with 100 μl osmotic mini pumps (Alzet) implanted at the same stereotaxic coordinates as above.

### Reversal learning

Spatial memory was assessed by the Morris water maze test (D’Hooge & Deyn, 2001) in which mice use visual cues to locate an escape platform (9 cm in diameter) in an open circular swimming arena (150 cm in diameter, 40 cm deep) filled with opaque water (Acusol, 20 ± 1 °C). The escape platform was hidden 1 cm below the water surface. Room temperature was kept constant at 24°C and both arena placement and surrounding visual cues were kept fixed during all experiments. Data was acquired by the SMART recording system and tracking software (Panlab). Data processing was automated with NAT (Navigation Analysis Tool), an in-house tool rooted in MATLAB (Jarlier *et al*, 2013). Mice underwent a two-phase training protocol (Fig. 4*E*). The first phase consisted of 5 training days, 1 day of rest followed by probe test 1 (initial learning). For the second phase, the platform was moved to a different quadrant, and again training lasted 5 days followed by 1 day of rest before probe test 2 (reversal learning). Training sessions consisted of 4 daily swimming trials (1 h interval between trials) starting randomly from different positions, with each quadrant sampled once per day. For each trial, mice were released at a starting point facing the inner wall and given a maximum of 90 s to locate and climb onto the escape platform. Mice that failed to locate the platform within 90 s were guided to it. In either case, mice were allowed to stay on the platform for 30 s. To assess spatial memory, probe tests were performed 24 h after the last training session by tracking mice for 60 s with the platform absent. Measured parameters during the training phases were the average time taken to reach the platform (i.e., mean escape latency) and the average speed. Measured parameters during probe tests were the mean distance traveled by the mouse before arriving at the platform location, % time spent in the target quadrant, and the number of platform location crossings.

### Electrophysiology

Acute transverse hippocampal slices (400 μm) were prepared as previously described (Pannasch *et al*, 2014). Briefly, mice were culled by cervical dislocation and decapitation. Brains were rapidly removed and placed in chilled (1-4 °C) artificial cerebrospinal fluid (aCSF) composed of (in mM) 119 NaCl, 2.5 KCl, 0.5 CaCl2, 1.3 MgSO4, 1 NaH2PO4, 26.2 NaHCO3 and 11 glucose. After sectioning, hippocampal slices were maintained at room temperature in a storage chamber containing aCSF saturated with 95% O2 and 5% CO2 for at least 1 h before the experiments. Slices were then transferred to a submerged recording chamber mounted on a Scientifica SliceScope Pro 6000 microscope equipped for infrared-differential interference (IR-DIC) microscopy and were perfused with aCSF (2 ml/min) at RT. All experiments were performed in CA1 stratum radiatum region of the hippocampus. Field excitatory postsynaptic potentials (fEPSPs) were recorded with glass pipettes (2–5 MΩ) filled with 1 M NaCl. Postsynaptic responses were evoked by stimulating Schaffer collaterals (0.033 Hz) in CA1 stratum radiatum with aCSF-filled glass pipettes. Input/output relationships of evoked excitatory postsynaptic potentials (EPSPs) were assayed by incrementing stimulation strength (5 to 50 μA, 100 μs). The test-shock used in subsequent experiments was chosen to elicit 50% of the maximal slope. Paired-pulse experiments consisted of 2 identical stimuli with increasing inter-pulse intervals (50 to 250 ms). Paired-pulse ratios were generated by plotting the maximum slope of the second fEPSP as a percentage of the first. Long-term potentiation (LTP) was induced by high-frequency stimulation (HFS: 2 trains of 100 pulses at 100 Hz, 30 s intertrain interval).

### PCR and Western blots

Mice were anesthetized for intracardiac perfusion with PBS. Subsequently, choroid plexus, hippocampus, cortex, retina and/or SVZ samples were extracted in ice-cold PBS, frozen on dry ice, and stored at −20 °C. Genomic DNA, total RNA and proteins were recovered by using the Allprep DNA/RNA/Protein Mini Kit (Qiagen 80004). The efficiency of Cre-induced recombination in *App_flox/flox_* mice was verified by PCR with primers F, C and D previously described (Mallm *et al*, 2010). For quantitative RT-PCR, cDNA was synthetized from 13 ng total RNA with QuantiTect Reverse Transciption Kit (Qiagen 205313). Quantitative PCR reactions were carried out in triplicate with SYBR Green I Master Mix (Roche S-7563) on a LightCycler 480 system (Roche). Expression was calculated by using the *2^-ΔΔCt^* method with *Gapdh* as a reference. For Western blot analysis, proteins samples were separated on NuPAGE 4-12% Bis-Tris pre-cast gels (Invitrogen NP0321) at 200 V for 1 h and transferred onto a methanol-activated PVDF membrane at 400 mV for 1 h. Primary antibody anti-APP (22C11 mouse 1:500, Millipore MAB348) was incubated overnight at 4 °C and anti-mouse-HRP-coupled secondary antibody (1:3000, Life Technologies) was incubated for 1.5 h at RT. Signal was detected by SuperSignal West Femto Substrate (Thermo Scientific 34095) with a LAS-4000 gel imager (Fujifilm) and quantified by densitometry with ImageJ.

### Histology

Mice were anesthetized for intracardiac perfusion with PBS, and brains were subsequently removed and fixed in 4% paraformaldehyde for 3 days. Coronal sections (60 μm) were obtained by Vibratome (Microm). For immunostaining, floating sections were steamed 10 min in citrate buffer (10 mM sodium citrate pH 6.0, 0.05% Tween), washed in PBS and treated with blocking solution (PBS pH 7, 1% triton-X100, 5% normal goat serum) for 1 h. Primary antibody anti-Ki67 (SP6 rabbit 1:500, Abcam ab16667) was incubated overnight at 4 °C in blocking solution and biotinylated anti-rabbit IgG secondary antibody (1:3000, Vector Laboratories BA-1000) at RT for 1h. Staining was visualized by using the Vectastain ABC HRP kit (Vector Laboratories PK-6100) with the DAB Peroxidase (HRP) Substrate Kit (Vector Laboratories SK-4100). Ki67-positive cells were counted manually in the DG on 5 sections per mouse (sections 180 μm apart, n = 5 to 8 mice per group). Cells in SVZ were counted by StereoInvestigator (Stereology Software, MBF Bioscience) on 8 sections per mouse (sections 180 μm apart, n = 5 to 8 mice per group).

### Statistical Analysis

Morris water maze training data (escape latency and speed) were analyzed with two-way repeated measures ANOVA with Statview 5.0.1 (SAS). Probe test data were analyzed for normality (D’Agostino & Pearson omnibus normality test) and unpaired *t* test was used for group comparisons with Statview 5.0.1 (SAS). Electrophysiological data were analyzed by ANOVA with Statistica 6.1 and Statview 5.0.1. Histological and biochemical data were analyzed by unpaired *t* test or ANOVA (as described in Figure legends) with Prism v6 (GraphPad). All data are given as mean ± standard error of the mean (SEM).

## Acknowledgments

The authors thank Julien Schmitt for technical support. Funding was provided by ERC Advanced Grant HOMEOSIGN n°339379 (to AP) and by ANR-10-LABX-BioPsy (to LR-R).

## Author contributions

KA, VOM and AAD performed animal surgery, biochemistry, and histology experiments and analyzed data. JV and LR-R performed and analyzed behavior experiments. VOM and GD performed and analyzed electrophysiology experiments. CLP performed histological analyses. MR and UCM provided mouse models. KA, VOM, AP and AAD designed experiments. VOM, UCM, AP and AAD wrote the manuscript.

## The paper explained

### Problem

APP metabolites in cerebrospinal fluid (CSF) are biomarkers of Alzheimer disease progression and are assumed to originate from brain parenchyma waste. However, high APP expression in choroid plexus, which generates CSF, suggests CSF-borne APP metabolites have functional purpose.

### Results

Through genetic and viral approaches selective for adult mouse choroid plexus, we find that sAPPα ectodomain levels in CSF modulate neurogenic niche proliferation (both hippocampus and subventricular zone). Furthermore, by favoring Aβ peptide production in wild type adult mouse choroid plexus, we observe reduced niche proliferation, deficits in hippocampus synaptic plasticity, and behavioral defects related to learning.

### Impact

These findings highlight the potential for choroid plexus APP to regulate brain function and open a novel way for countering disease-induced impairments.

---

**Figure S1.**
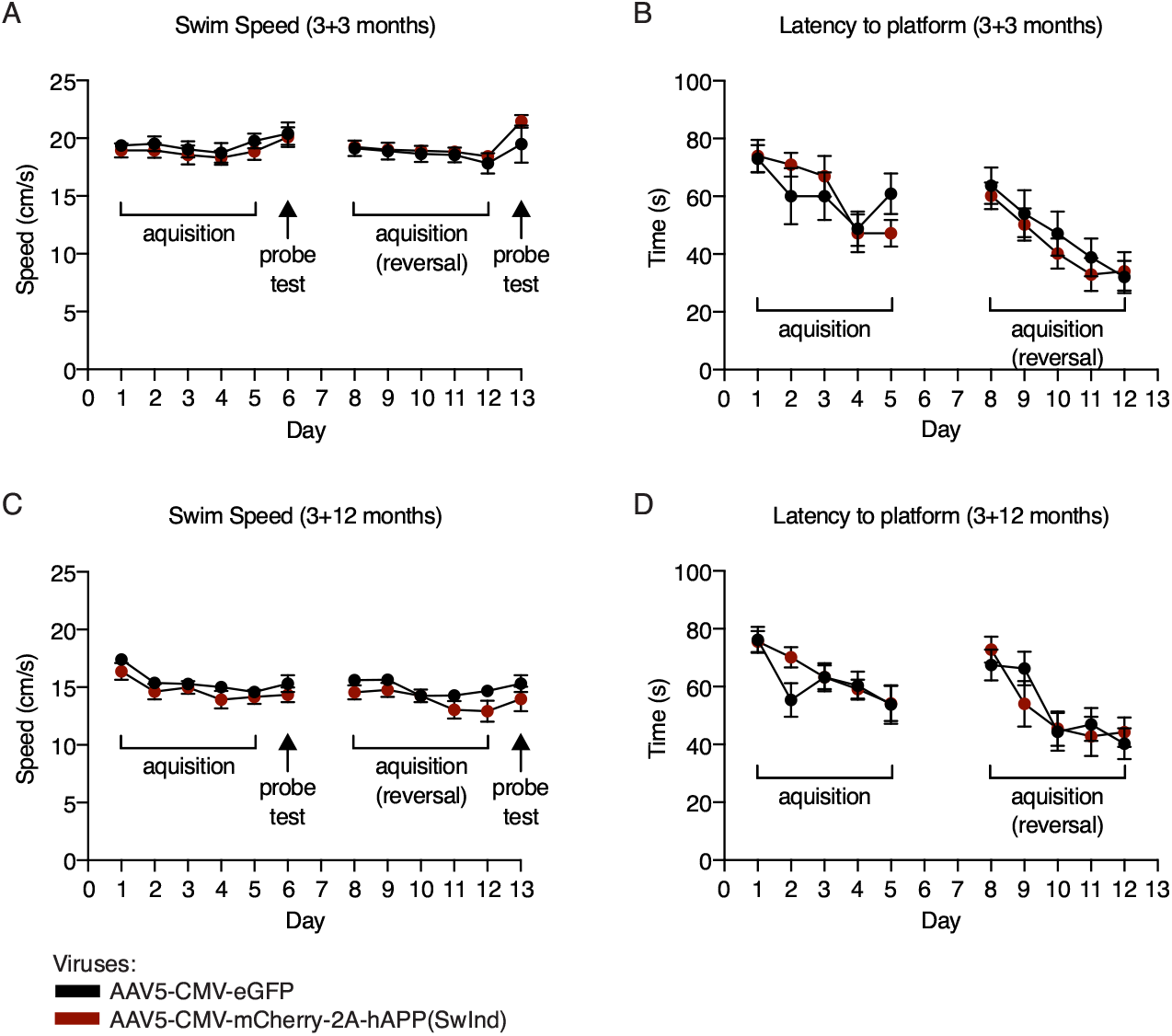
Morris water maze escape latency and speed measured during acquisition phases and probe tests. At 3-months **(A, B)** and 12-months **(C, D)** post viral injection, recordings were performed during the 5 days of initial and reversal learning phase to determine displacement speed (**A, C**) and mean escape latency (**B, D**). Displacement speed was also measured during probe tests (**A, C**). Escape latency cannot be measured during probe tests owing to the absence of a platform. Two-way repeated measures ANOVA; all values ±SEM.

